# L-selectin shedding regulates post-infarction neutrophilia without altering myocardial injury or ventricular remodelling

**DOI:** 10.64898/2026.11.12.765854

**Authors:** Davide Maselli, Samuel J Hart, Yue Yang, Mateusz Tomczyk, Silvia Trevelin, Diana Cash, Camilla Simmons, Greta Sawyer, Xiaohong Zhang, Mauro Giacca, Ajay Shah, Aleksandar Ivetic

## Abstract

Myocardial infarction (MI) triggers rapid neutrophil mobilisation from the bone marrow (BM), resulting in systemic neutrophilia that is widely associated with myocardial injury. However, whether the magnitude of this response directly influences cardiac injury and subsequent ventricular remodelling remains unclear. Ectodomain shedding of L-selectin is a hallmark of neutrophil activation, although its contribution to neutrophil mobilisation following MI has not been established. Using mice expressing a non-cleavable L-selectin mutant (L(E)), we investigated the role of L-selectin shedding in post-MI neutrophilia. Under homeostatic conditions, WT and L(E) mice exhibited comparable BM neutrophil maturation and circulating neutrophil numbers. Following permanent coronary ligation, however, L(E) mice exhibited markedly attenuated neutrophilia, associated with reduced circulating CXCL2 and decreased surface CXCR2 expression on BM neutrophils. Intravenous recombinant CXCL2 increased circulating neutrophil numbers in WT but not L(E) mice, suggesting that impaired mobilisation cannot be explained solely by reduced receptor/ligand availability. Despite attenuated neutrophilia, myocardial neutrophil number was comparable between genotypes at 24 hours post-MI. Circulating cardiac troponin I concentrations and MRI-quantified infarct size also showed no detectable differences. Longitudinal cardiac MRI at days 1, 14 and 28 revealed comparable ventricular structure and function, while histological scar size was similar at day 28. These findings identify L-selectin shedding as a regulator of post-MI neutrophilia and suggest that the magnitude of this systemic response can be dissociated from acute myocardial injury and subsequent ventricular remodelling.

## Introduction

Myocardial infarction (MI) triggers a rapid inflammatory response characterised by the mobilisation of neutrophils from the bone marrow (BM), resulting in systemic neutrophilia and subsequent infiltration of the infarcted myocardium^1^. This response is conserved between mice and humans and represents an important component of the early inflammatory phase following MI. Neutrophils can exacerbate myocardial injury through the generation of reactive oxygen species (ROS), release of cytotoxic enzymes and amplification of local inflammatory signalling^2–4^. However, neutrophils also contribute to the clearance of necrotic tissue and subsequent inflammatory resolution, highlighting their complex involvement in myocardial repair^5^. Although elevated circulating neutrophil numbers are associated with myocardial injury, whether the magnitude of systemic neutrophilia directly determines myocardial neutrophil accumulation, acute injury or subsequent ventricular remodelling remains unclear.

Neutrophil mobilisation from the BM is regulated by the balance between retention signals mediated principally through CXCR4/CXCL12 and mobilisation signals involving CXCR2 and its chemokine ligands, including CXCL1 and CXCL2^6–8^. Following inflammatory stimulation, CXCR2-dependent signalling promotes neutrophil release into the circulation, contributing to systemic neutrophilia. Neutrophils themselves can produce CXCL2, supporting autocrine and paracrine signalling that amplifies inflammatory recruitment^9^. Furthermore, recent evidence indicates that neutrophils recruited to the ischaemic myocardium can originate from vascular marginated populations, suggesting that myocardial recruitment is not necessarily dependent on newly mobilised BM neutrophils^1^. Nevertheless, the molecular mechanisms regulating neutrophil release from the BM following MI remain incompletely understood.

L-selectin (CD62L) is a type I transmembrane adhesion receptor expressed on most circulating leukocytes, including neutrophils, and contributes to leukocyte tethering, rolling and recruitment across the vascular endothelium^10,11^. Following neutrophil activation, L-selectin undergoes rapid proteolytic ectodomain shedding, predominantly mediated by ADAM17, resulting in the release of soluble L-selectin and reduced surface receptor expression^12–14^. Although shedding is widely recognised as a hallmark of neutrophil activation, its contribution to neutrophil mobilisation remains poorly understood. Studies using mice expressing a non-cleavable form of L-selectin have demonstrated altered leukocyte recruitment during experimental inflammation, indicating that ectodomain shedding regulates trafficking^15,16^.

The potential relevance of L-selectin shedding to MI is supported by clinical observations demonstrating an inverse association between neutrophil surface L-selectin expression and circulating neutrophil numbers during inflammation^17^. In patients with acute MI, reduced neutrophil surface L-selectin expression has also been associated with increased neutrophilia and larger infarct size^18^. Earlier studies reported changes in circulating soluble L-selectin following MI, although these observations did not establish whether reduced surface expression resulted directly from ectodomain shedding^19,20^. More recently, experimental MI has been associated with reduced neutrophil surface L-selectin and increased circulating soluble L-selectin, consistent with enhanced receptor shedding during the acute inflammatory response^1^. Nevertheless, whether L-selectin shedding contributes directly to post-infarction neutrophil mobilisation, or merely accompanies neutrophil activation, remains unresolved.

Here, we investigated the contribution of L-selectin shedding to the systemic neutrophilic response following permanent coronary artery ligation using mice expressing a non-cleavable L-selectin mutant (L(E))^15^. We examined the relationship between L-selectin shedding, CXCL2/CXCR2-associated mobilisation and post-infarction neutrophilia. We further asked whether attenuation of this response was accompanied by changes in myocardial neutrophil numbers, acute myocardial injury or subsequent ventricular remodelling. By combining analysis of neutrophil number with longitudinal cardiac magnetic resonance imaging and histological assessment of scar formation, we sought to distinguish the regulation of systemic neutrophilia from its potential contribution to myocardial injury and repair.

## Results

### L-selectin shedding does not affect steady-state neutrophil abundance or bone marrow maturation

Previous studies using mice expressing a non-cleavable L-selectin mutant (L(E)) reported comparable circulating and bone marrow (BM) neutrophil populations to those of wild-type (WT) littermates^15^. We confirmed these observations in L(E) mice maintained on a C57BL/6 background, with no significant differences in the proportion of neutrophils among BM or circulating CD45⁺ leukocytes (**Fig. 1A,B**). Although surface L-selectin expression was comparable between genotypes on BM neutrophils, circulating L(E) neutrophils exhibited significantly higher L-selectin expression than their WT counterparts (**Fig. 1C,D**), consistent with impaired receptor shedding.

**Figure 1.**
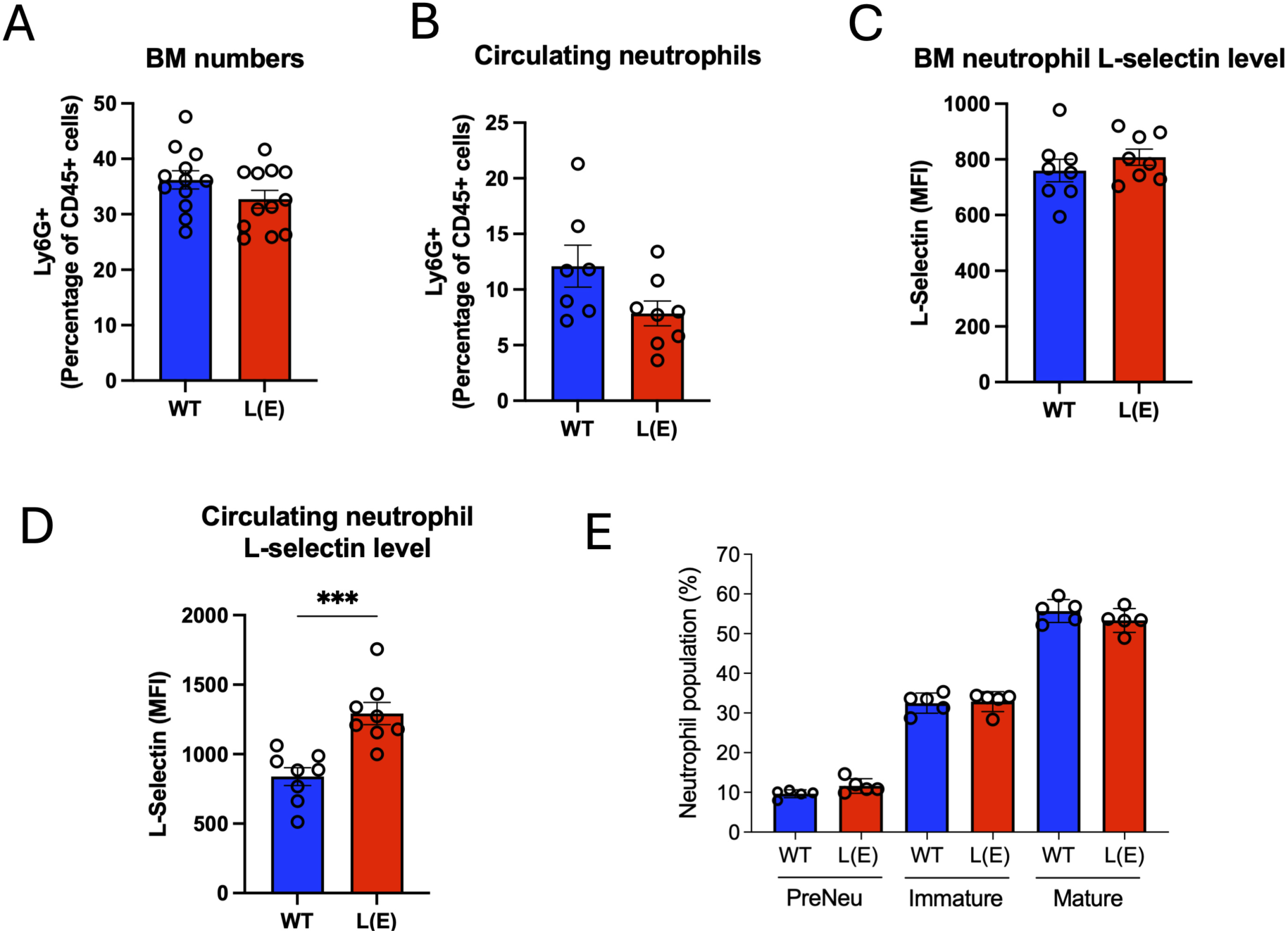
Neutrophil numbers and bone marrow maturation are unaffected in naive L(E) mice, which carry elevated surface L-selectin on circulating neutrophils. (**A**) Bone marrow (BM) neutrophils in naive WT and L(E) mice, identified by flow cytometry as Ly6G⁺ events and expressed as a percentage of CD45⁺ leukocytes. (**B**) Circulating neutrophils in the same animals, expressed as a percentage of CD45⁺ leukocytes. (**C**) Surface L-selectin on BM neutrophils, expressed as median fluorescence intensity (MFI). (**D**) Surface L-selectin on circulating neutrophils (MFI). (**E**) Distribution of BM neutrophil maturation subsets, gated as described in Methods (taken from Evrard et al^24^) and expressed as a percentage of total BM neutrophils: CD117ʰⁱCXCR4ʰⁱ pre-neutrophils (PreNeu), Ly6G⁻ᐟᵒCXCR2⁻ᐟᵒ immature neutrophils and Ly6G⁺CXCR2⁺ mature neutrophils. Bars show mean ± SEM with each symbol representing one mouse (n = 13–14 per genotype in A; n = 7–8 in B–D; n = 5 in E). Blue, WT; red, L(E). Unpaired t-test (A–D); two-way ANOVA with Šidák’s multiple comparisons test (E). ***p < 0.001; comparisons without an asterisk were not significant.

L-selectin is expressed during early haematopoietic differentiation and has been identified on lymphoid-primed progenitor populations^21–23^. However, whether preventing L-selectin shedding influences neutrophil maturation within the BM has not been established. We therefore examined neutrophil maturation in naïve WT and L(E) mice using the flow cytometric gating strategy described by Evrard and colleagues^24^. Following exclusion of CD170⁺ eosinophils and CD115⁺ monocytes, neutrophils were identified within the Gr1⁺CD11b⁺ population and classified into CD117ʰⁱCXCR4ʰⁱ pre-neutrophils (PreNeu), Ly6GˡᵒCXCR2ˡᵒ immature neutrophils and Ly6G⁺CXCR2⁺ mature neutrophils. The relative proportions of these three populations were comparable between WT and L(E) mice (**Fig. 1E**). Together, these findings indicate that preventing L-selectin shedding does not detectably alter steady-state neutrophil numbers or the distribution of BM neutrophil maturation subsets, providing a baseline for investigating its contribution to inflammatory neutrophil mobilisation.

### L(E) neutrophils retain chemoattractant responsiveness despite impaired L-selectin shedding

Much of our understanding of L-selectin-dependent neutrophil trafficking has been derived from studies using L-selectin-deficient mice, which established important roles for this adhesion receptor in leukocyte rolling and recruitment during inflammation^25–28^. However, complete deletion of L-selectin cannot distinguish the adhesive functions of the intact receptor from those regulated specifically by ectodomain shedding. Studies using L(E) mice have demonstrated that preventing L-selectin shedding alters neutrophil recruitment during thioglycollate-induced peritonitis and impairs neutrophil responses during bacterial pneumonia^15,16^. Nevertheless, comparatively little is known about the contribution of L-selectin shedding to neutrophil trafficking in response to defined inflammatory chemoattractants.

We first investigated neutrophil recruitment into the peritoneal cavity following intraperitoneal administration of CXCL2. Four hours after challenge, the proportion of neutrophils among recovered peritoneal leukocytes was comparable between WT and L(E) mice (**Fig. 2A**). However, whereas WT neutrophils recovered from the peritoneum exhibited significantly reduced surface L-selectin expression relative to circulating neutrophils, L(E) neutrophils retained substantially higher surface expression (**Fig. 2B**). These findings indicate that the L(E) mutation does not detectably alter neutrophil numbers in this acute CXCL2-induced inflammatory model, despite differences in surface L-selectin regulation.

**Figure 2.**
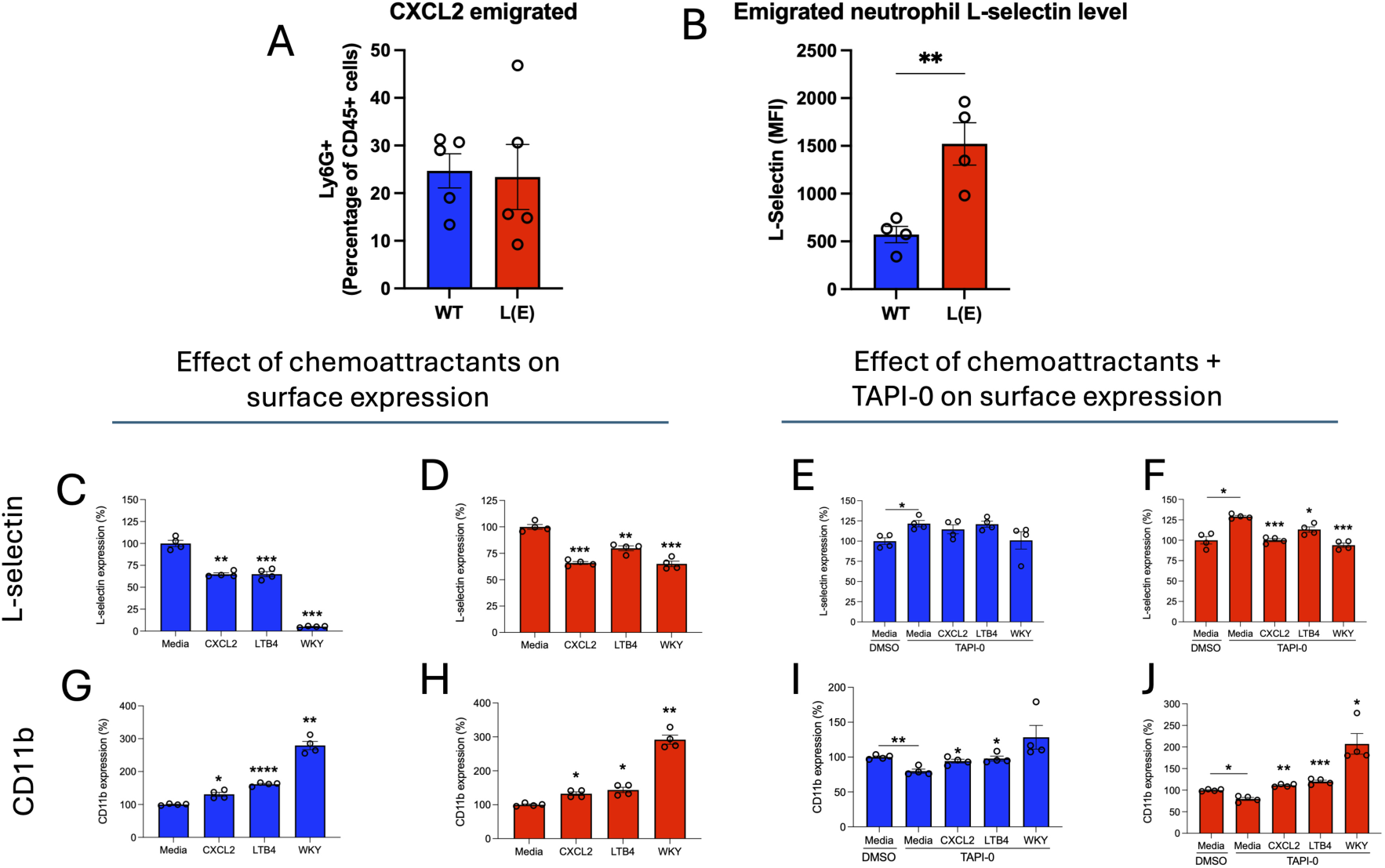
The L(E) mutation does not impair neutrophil trafficking to a CXCL2-challenged peritoneum and does not prevent Mac-1 upregulation in response to chemoattractant stimulation. (**A**) Neutrophils recovered by peritoneal lavage 4 h after intraperitoneal CXCL2, expressed as a percentage of CD45⁺ leukocytes. Neutrophils identified as Ly6G^+^, CD11b^+^ (**B**) Surface L-selectin (MFI) on emigrated peritoneal neutrophils. (**C**, **D**) BM-derived neutrophils from WT (**C**) and L(E) (**D**) mice were stimulated for 30 min with medium alone (RPMI, FCS, pen/strep), CXCL2, LTB₄ or the formyl peptide WKYMVm (WKY; all at 500 nM), and surface L-selectin measured by flow cytometry; data are normalised to the unstimulated (media) control of the same genotype. (**B**, **F**) As in (**C**) and (**D**), with cells pre-incubated for 30 min with the metalloproteinase inhibitor TAPI-0 (10 µM) or an equivalent volume of DMSO before stimulation; WT neutrophils are shown in **E** and L(E) neutrophils in **F**. (**G**, **H**) Surface CD11b (Mac-1) on WT (**G**) and L(E) (**H**) neutrophils from the experiments in **C** and **D**, normalised to the media control of the same genotype. (**I**, **J**) Surface CD11b on WT (**I**) and L(E) (**J**) neutrophils from the TAPI-0 experiments in (**E**) and (**F**). Bars show mean ± SEM, with each symbol representing an independent experiment (n = 4 in **C**–**J**). Blue, WT; red, L(E). In **C**, **D**, **G** and **H**, comparisons are against the media control of the same genotype. In **E**, **F**, **I** and **J**, all comparisons -including that for the DMSO-treated media control -are against the TAPI-0-treated media control. Welch’s ANOVA with Dunnett’s T3 multiple comparisons test throughout. *p < 0.05; **p < 0.01; ***p < 0.001; ****p < 0.0001.

To investigate the effects of chemoattractant stimulation directly, BM-derived neutrophils were challenged with CXCL2, leukotriene B4 (LTB4) or the formyl peptide WKYMVm. All three agonists induced reductions in surface L-selectin expression, although the magnitude of this response differed between genotypes and stimuli (**Fig. 2C,D**). CXCL2 and LTB4 induced relatively modest reductions in both WT and L(E) neutrophils, whereas WKYMVm produced a pronounced loss of surface L-selectin in WT cells that was substantially attenuated in L(E) neutrophils. Notably, L(E) neutrophils nevertheless exhibited an approximately 30–35% reduction in surface L-selectin following stimulation, indicating that the mutation does not completely prevent agonist-induced loss of receptor expression.

We next examined whether these reductions were sensitive to metalloproteinase inhibition. Pretreatment with the broad-spectrum metalloproteinase inhibitor TAPI-0 largely prevented the loss of surface L-selectin in both WT and L(E) neutrophils (**Fig. 2E,F**). These observations suggest that the residual reduction in surface L-selectin on L(E) neutrophils involves a metalloproteinase-sensitive process, despite disruption of the canonical membrane-proximal cleavage region. Whether this reflects proteolysis at an alternative site or another mechanism affecting surface receptor availability remains to be determined.

L-selectin shedding and upregulation of the β2 integrin Mac-1 (CD11b/CD18; αMβ2) are closely associated features of neutrophil activation^10^. Previous studies have also demonstrated that L-selectin-dependent signalling can influence Mac-1 expression and function^29^. We therefore examined whether impaired L-selectin shedding in L(E) neutrophils affected chemoattractant-induced CD11b upregulation.

In WT neutrophils, CXCL2 and LTB4 induced relatively modest increases in surface CD11b, whereas WKYMVm elicited a more pronounced response (**Fig. 2G**). A similar pattern was observed in L(E) neutrophils, including substantial CD11b upregulation following WKYMVm stimulation despite the markedly reduced loss of surface L-selectin (**Fig. 2H**). Furthermore, TAPI-0 pretreatment had comparatively modest effects on agonist-induced CD11b expression in either genotype (**Fig. 2I,J**). Collectively, these findings demonstrate that L(E) neutrophils retain the capacity to respond to inflammatory chemoattractants through CD11b upregulation, even when L-selectin shedding is substantially impaired. Thus, the L(E) mutation does not produce a generalised defect in the activation responses examined here, allowing its contribution to post-infarction neutrophil mobilisation to be investigated independently of these responses.

### L-selectin shedding regulates post-infarction neutrophilia and bone marrow CXCR2 expression

Neutrophilia is a characteristic feature of the early inflammatory response following myocardial infarction (MI), reflecting the rapid mobilisation of neutrophils from the BM into the circulation^1^. Although WT and L(E) mice exhibited comparable circulating neutrophil proportions under homeostatic conditions (**Fig. 1B**), a marked difference emerged following permanent ligation of the left anterior descending (LAD) coronary artery. At 24 h post-MI, WT mice exhibited a pronounced increase in circulating neutrophil numbers, whereas this response was substantially attenuated in L(E) mice (**Fig. 3A**). These findings suggest that L-selectin shedding contributes to the systemic neutrophilic response following MI.

**Figure 3.**
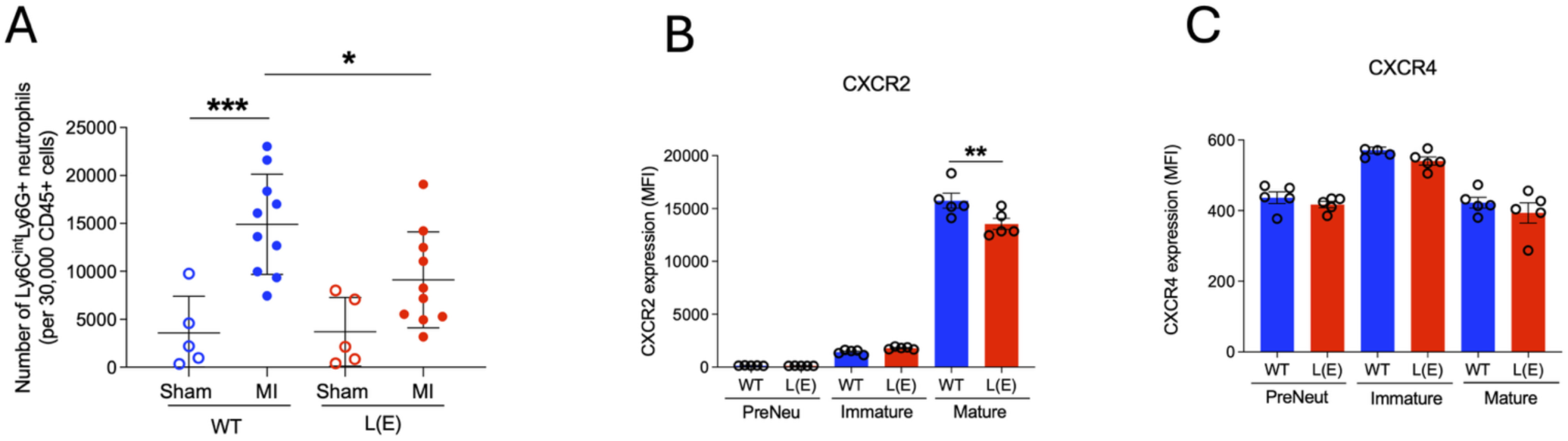
L(E) mice fail to mount neutrophilia after myocardial infarction and show reduced CXCR2 on mature bone marrow neutrophils. (**A**) Circulating neutrophils (Ly6CⁱⁿᵗLy6G⁺ events per 30,000 CD45⁺ cells) in WT and L(E) mice 24 h after sham surgery or permanent LAD ligation (MI). (**B**) Surface CXCR2 (MFI) on BM pre-neutrophil (PreNeu), immature and mature neutrophil subsets of naive WT and L(E) mice. (**C**) Surface CXCR4 (MFI) on the same BM subsets. Data are mean ± SD (**A**) or mean ± SEM (**B**, **C**), with each symbol representing one mouse (n = 4–5 sham and n = 10 MI in **A**; n = 5 per genotype in **B** and **C**). Blue, WT; red, L(E). Two-way ANOVA with Šidák’s multiple comparisons test. *p < 0.05; **p < 0.01; ***p < 0.001.

The mobilisation of neutrophils from the BM is regulated by opposing chemokine signals, with CXCL12/CXCR4 promoting retention and CXCL2/CXCR2 signalling facilitating their release into the circulation^6–8^. We therefore examined whether the impaired neutrophilic response in L(E) mice was associated with altered expression of these chemokine receptors. Flow cytometric analysis of BM neutrophil maturation subsets revealed significantly reduced surface CXCR2 expression on mature L(E) neutrophils compared with WT counterparts (**Fig. 3B**). In contrast, surface CXCR4 expression was comparable between genotypes across the BM neutrophil populations examined (**Fig. 3C**). These findings identify reduced CXCR2 surface expression as a potential contributor to the attenuated neutrophilic response in L(E) mice, without evidence of a corresponding increase in CXCR4 expression. Together, the results suggest that L-selectin shedding influences the mechanisms governing inflammatory neutrophil mobilisation following MI.

### Attenuated post-infarction neutrophilia in L(E) mice is associated with reduced circulating CXCL2

Having identified reduced CXCR2 expression on mature BM neutrophils in L(E) mice, we next investigated whether the availability of chemokine ligands involved in neutrophil mobilisation was also altered. CXCR2 signalling, particularly through CXCL1 and CXCL2, promotes neutrophil release from the BM^6,7^. Furthermore, neutrophils themselves can produce CXCL2, contributing to the amplification of inflammatory recruitment through autocrine and paracrine signalling^9^. We therefore examined whether preventing L-selectin shedding was associated with changes in circulating inflammatory mediators following MI.

Plasma samples collected from WT and L(E) mice at days 1 and 5 following permanent LAD ligation were analysed using an antibody array comprising 144 analytes, including cytokines, chemokines, growth factors, soluble receptors, adhesion molecules and proteases (see Methods). Of these, 28 analytes exhibited nominal genotype-associated differences (p < 0.05), all displaying lower signal intensities in L(E) mice (**Fig. 4A**). Among the analytes identified, soluble L-selectin exhibited reduced circulating levels in L(E) mice (**Fig. 4A**), consistent with the shedding-resistant phenotype previously described in this model^15^. Notably, CXCL2 (MIP-2) exhibited the largest genotype-associated difference, with plasma signal intensities approximately 3.43-fold lower in L(E) mice than WT mice at day 1 post-MI and 1.66-fold lower at day 5 (**Fig. 4B**). In the same array dataset, comparisons of CXCL2 at individual timepoints yielded nominal p values of 0.0081 and 0.0279, respectively.

**Figure 4.**
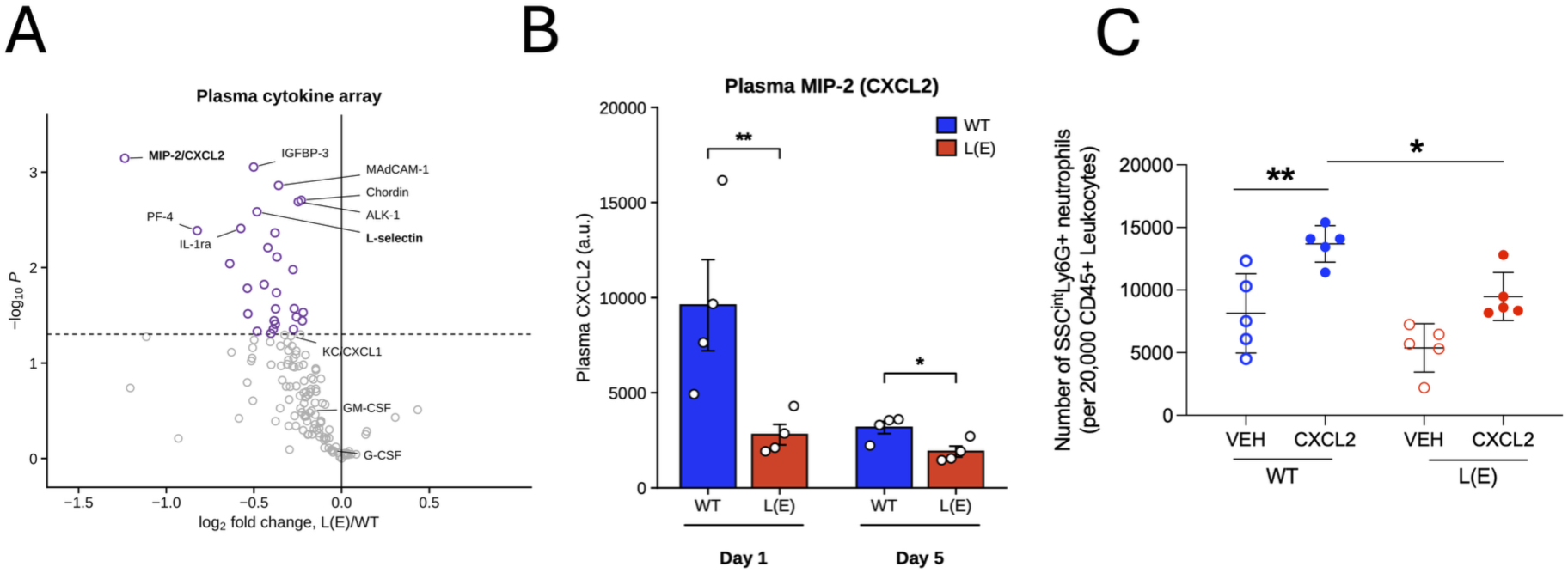
Reduced circulating CXCL2 following myocardial infarction is associated with impaired CXCL2-induced neutrophilia in L(E) mice. (**A**) Volcano plot of 144 plasma analytes measured using a cytokine antibody array in wild-type (WT) and L-selectin shedding-resistant (L(E)) mice at days 1 and 5 following permanent left anterior descending (LAD) coronary artery ligation (n = 4 mice per genotype per timepoint). The x-axis represents the log₂ fold change (L(E)/WT), and the y-axis represents the −log₁₀ p value derived from two-way ANOVA of log₂-transformed signal intensities, with genotype and timepoint as fixed factors. The dashed line indicates p = 0.05. Purple symbols identify 28 analytes exhibiting nominal genotype-associated differences (p < 0.05), all of which were lower in L(E) mice; grey symbols indicate analytes not reaching this threshold. (**B**) Plasma CXCL2 (MIP-2) signal intensities in WT and L(E) mice at days 1 and 5 post-MI, determined from the same antibody array. CXCL2 was 3.43-fold lower in L(E) mice at day 1 and 1.66-fold lower at day 5. Data are mean ± SEM, with each symbol representing one mouse (n = 4 per genotype per timepoint). Unpaired Student’s t-tests were performed on log₂-transformed intensities at each timepoint (day 1, p = 0.0081; day 5, p = 0.0279). (**C**) Acute systemic administration of recombinant murine CXCL2 in non-infarcted male WT and L(E) mice. Mice received recombinant CXCL2 (1 µg in 100 µL phosphate-buffered saline (PBS)) or PBS alone (vehicle) by intravenous tail-vein injection. After 45 min, circulating leukocytes were isolated and stained for flow cytometric analysis. Neutrophils were identified as CD45⁺SSCⁱⁿᵗLy6G⁺ events and quantified within a fixed acquisition population of 30,000 CD45⁺ leukocytes. Data are mean ± SD (n = 5 mice per group). Statistical comparisons were performed using two-way ANOVA with Šidák’s multiple comparisons test (*p ≤ 0.05; **p ≤ 0.01). Recombinant CXCL2 induced a significant increase in circulating neutrophil numbers in WT mice, whereas the response in L(E) mice did not reach statistical significance.

In contrast, circulating G-CSF and GM-CSF, which contribute to granulopoiesis and neutrophil production, showed no detectable genotype-associated differences at either timepoint. Similarly, CXCL12 and CXCL1 did not exhibit statistically significant differences, although CXCL1 showed a trend towards lower expression in L(E) mice. The prominence of CXCL2 among the candidate mediators, together with reduced CXCR2 expression on mature BM neutrophils (**Fig. 3B**), suggested that altered CXCL2/CXCR2 signalling might contribute to the attenuated neutrophilic response observed in L(E) mice.

To investigate whether reduced circulating CXCL2 availability was sufficient to explain this phenotype, we administered recombinant murine CXCL2 intravenously to non-infarcted WT and L(E) mice. Within 45 min, WT mice exhibited an increase in the proportion of circulating neutrophils, whereas L(E) mice showed no comparable response (**Fig. 4C**). Thus, exogenous CXCL2 failed to elicit the neutrophilic response observed in WT mice, indicating that reduced circulating ligand availability alone is unlikely to account for the impaired response in L(E) animals.

Collectively, these findings demonstrate that preventing L-selectin shedding markedly attenuates the relative circulating neutrophilic response following MI. This phenotype is associated with reduced CXCR2 expression on mature BM neutrophils and lower circulating CXCL2 signals, while the failure of exogenous CXCL2 to elicit neutrophilia in L(E) mice suggests an additional impairment in the mechanisms governing neutrophil mobilisation. Together, these observations implicate L-selectin shedding in the regulation of the CXCL2/CXCR2-associated neutrophilic response following MI.

### Myocardial neutrophil number is preserved in L(E) mice despite attenuated post-infarction neutrophilia

Having established that L(E) mice exhibit markedly attenuated circulating neutrophilia following myocardial infarction (MI), we next investigated whether this response was accompanied by reduced neutrophil accumulation within the infarcted myocardium. Neutrophils rapidly infiltrate the ischaemic myocardium following coronary artery occlusion, where they contribute to the early inflammatory response^2,3^. However, recent evidence suggests that myocardial neutrophil recruitment can also involve pre-existing vascular marginated populations, raising the possibility that recruitment to the infarcted heart may not depend exclusively on newly mobilised bone marrow (BM) neutrophils^1^.

WT and L(E) mice underwent permanent ligation of the left anterior descending (LAD) coronary artery, and hearts were harvested 24 h later. The myocardium was separated into basal and apical regions, corresponding respectively to tissue above and below the ligation site. Following enzymatic digestion, recovered leukocytes were analysed by flow cytometry to determine the number of neutrophils within the CD45⁺ population. In sham-operated mice, neutrophil number was low in both myocardial regions, with no detectable differences between genotypes (**Fig. 5A**). Following MI, neutrophil number increased markedly within the apical region of both WT and L(E) hearts (**Fig. 5B**). Despite the pronounced attenuation of circulating neutrophilia in L(E) mice (**Fig. 3A**), the proportion of neutrophils among recovered myocardial CD45⁺ leukocytes were comparable between genotypes in both the basal and apical regions (**Fig. 5B**).

**Figure 5.**
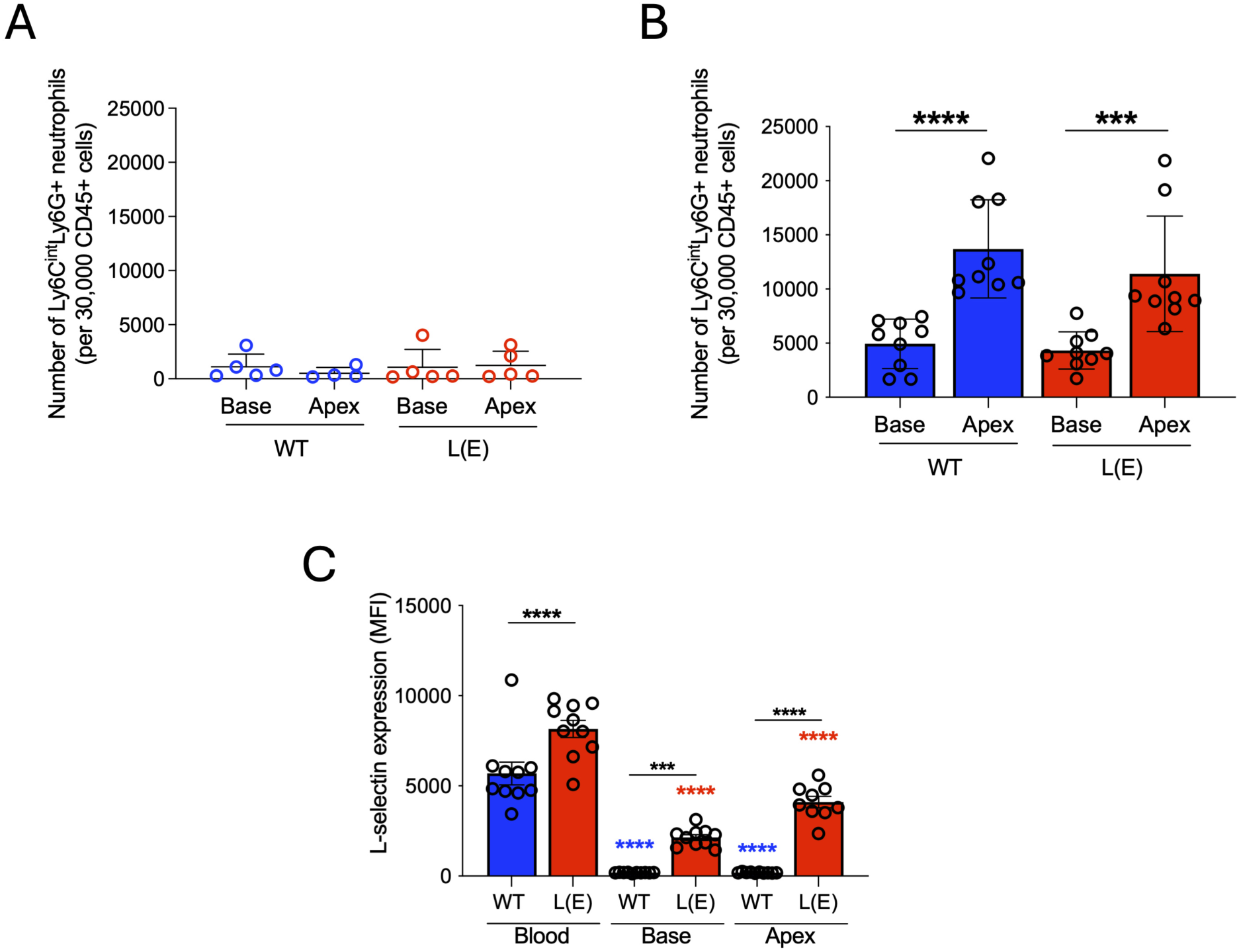
Myocardial neutrophil number is comparable between WT and L(E) mice despite attenuated circulating neutrophilia following MI. (**A**) Neutrophils (Ly6CⁱⁿᵗLy6G⁺ events per 30,000 CD45⁺ cells) recovered from the base and apex of WT and L(E) hearts 24 h after sham surgery. (**B**) Neutrophils recovered from the base and apex of WT and L(E) hearts 24 h after permanent LAD ligation. The apex corresponds to the infarcted region below the suture line and the base to the remote, non-infarcted myocardium. (**C**) Surface L-selectin (MFI) on neutrophils from blood, base and apex of the same infarcted animals. Black brackets indicate WT versus L(E) comparisons within a compartment; coloured asterisks indicate comparisons against circulating neutrophils of the same genotype (blue, WT; red, L(E)). Bars show mean ± SD with each symbol representing one mouse (n = 5 in A; n = 8–9 in B; n = 11–12 in C). Blue, WT; red, L(E). Two-way ANOVA with Šidák’s multiple comparisons test. ***p < 0.001; ****p < 0.0001.

We next examined surface L-selectin expression on neutrophils recovered from the infarcted myocardium. In WT mice, myocardial neutrophils exhibited markedly reduced surface L-selectin expression compared with circulating neutrophils, consistent with receptor downregulation during inflammatory recruitment (**Fig. 5C**). Although L(E) neutrophils also exhibited reduced surface L-selectin expression following myocardial recruitment, they retained significantly higher levels than their WT counterparts (**Fig. 5C**). This pattern was consistent with the partial reduction in surface L-selectin observed following chemoattractant stimulation of L(E) neutrophils in vitro (**Fig. 2C,D**).

Together, these findings demonstrate that the attenuated circulating neutrophilic response in L(E) mice is not accompanied by a detectable reduction in the relative number of neutrophils within the infarcted myocardium at 24 h.

### Attenuated post-infarction neutrophilia in L(E) mice is not accompanied by detectable differences in acute myocardial injury

Having established that L(E) mice exhibit markedly attenuated circulating neutrophilia following myocardial infarction (MI), despite comparable myocardial neutrophil number at 24 h (**Fig. 5**), we next investigated whether preventing L-selectin shedding influenced the extent of acute myocardial injury. Neutrophils contribute to the early inflammatory response following MI and have been implicated in myocardial injury through the generation of ROS, release of cytotoxic enzymes and amplification of local inflammation^2–4^. We therefore assessed myocardial injury using circulating cardiac troponin I (cTnI) concentrations and cardiac magnetic resonance imaging (MRI).

Plasma cTnI concentrations were measured 24 h following sham surgery or permanent ligation of the left anterior descending (LAD) coronary artery. MI induced a pronounced increase in circulating cTnI in both genotypes compared with their respective sham-operated controls (WT, 108.6 ± 11.5 ng/mL versus 5.5 ± 1.9 ng/mL; L(E), 86.1 ± 8.7 ng/mL versus 18.7 ± 7.0 ng/mL; p < 0.0001 for both comparisons; **Fig. 6A**). However, cTnI concentrations did not differ significantly between infarcted WT and L(E) mice (p = 0.147), indicating that the attenuated circulating neutrophilic response in L(E) mice was not accompanied by a detectable reduction in this biochemical marker of acute cardiomyocyte injury.

**Figure 6.**
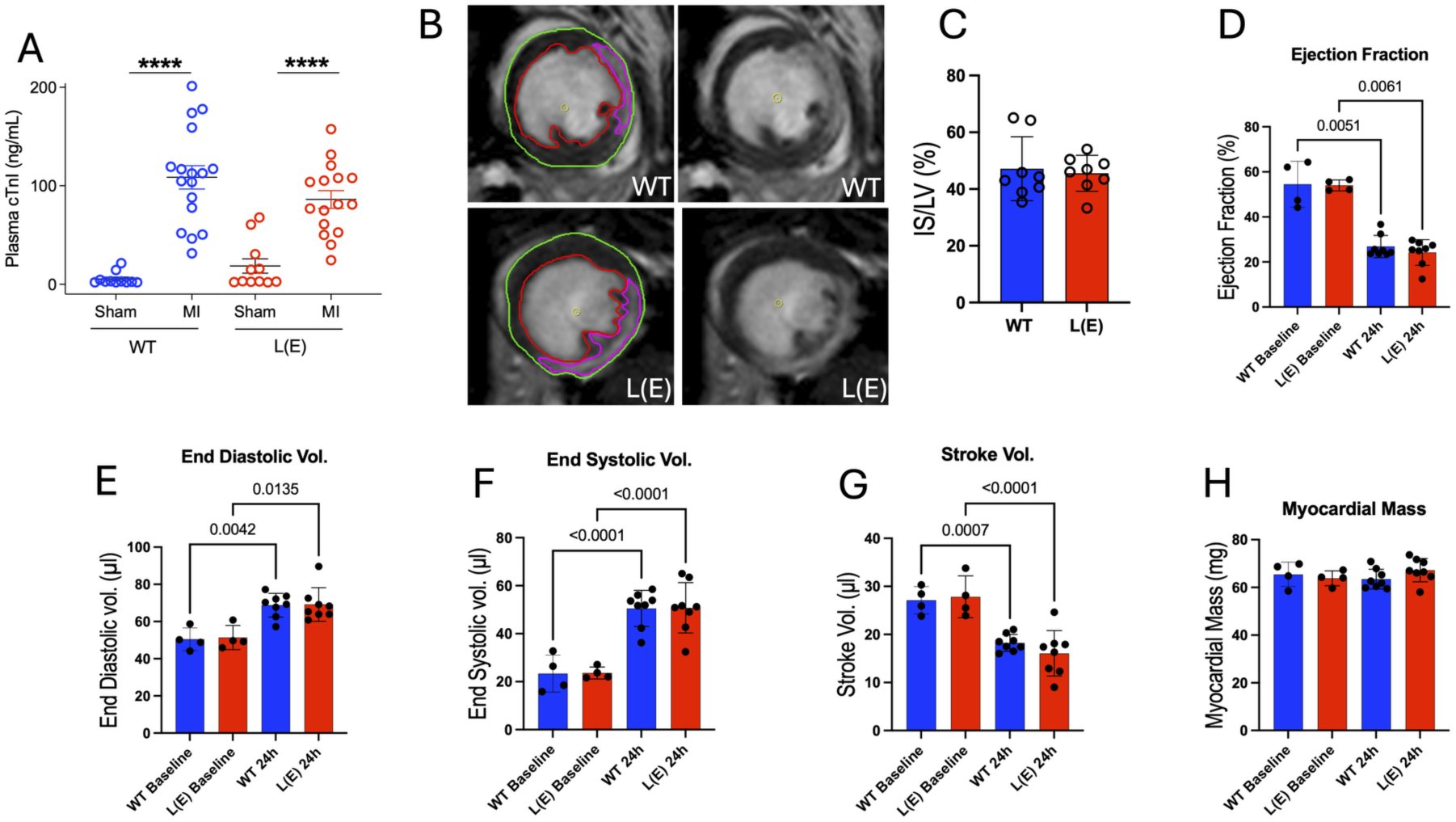
Attenuated post-infarction neutrophilia in L(E) mice is not accompanied by detectable differences in acute myocardial injury or ventricular function. (**A**) Plasma cardiac troponin I (cTnI) concentrations in wild-type (WT) and L-selectin shedding-resistant (L(E)) mice 24 h following sham surgery or permanent ligation of the left anterior descending (LAD) coronary artery. Data are presented as individual values (n = 11 sham and n = 16–17 MI mice per genotype). Statistical comparisons were performed using two-way ANOVA with Šidák’s multiple comparisons test. (**B**) Representative mid-ventricular short-axis late gadolinium-enhanced (LGE) cardiac magnetic resonance images acquired 24 h following MI in WT and L(E) mice, shown with (left) and without (right) segmentation contours. Green, epicardial boundary; red, endocardial boundary; magenta, region of late gadolinium enhancement. (**C**) Myocardial infarct size was quantified 24 h after permanent LAD ligation using late gadolinium enhancement (LGE) cardiac MRI, with infarct size defined using a threshold of two standard deviations above the mean signal intensity of remote myocardium. Infarct size was expressed as a percentage of left ventricular volume (IS/LV%). WT and L(E) mice (n = 8 per genotype) exhibited comparable infarct sizes. Data are presented as mean ± SEM. Statistical significance was assessed using a two-tailed unpaired Welch’s t-test (p = 0.731). (**D–H**) LV structural and functional parameters determined by cine cardiac MRI in WT and L(E) mice before (baseline) and 24 h after permanent LAD ligation: (**D**) LV ejection fraction (%); (**E**) LV end-diastolic volume (µL); (**F**) LV end-systolic volume (µL); (**G**) LV stroke volume (µL); and (**H**) LV myocardial mass (mg). Blue, WT; red, L(E). Bars represent mean ± SEM, with individual symbols representing separate animals. No statistically significant differences in circulating cTnI, MRI-derived infarct size or LV structural and functional parameters were detected between WT and L(E) mice following MI. ****p < 0.0001.

We next assessed infarct size using late gadolinium-enhanced (LGE) cardiac MRI at 24 h following permanent LAD ligation. Representative short-axis images demonstrated extensive enhancement within the anterolateral myocardium of both WT and L(E) mice, consistent with the expected distribution of injury following coronary artery occlusion (**Fig. 6B**). Infarct size was assessed using a semi-automated signal-intensity threshold set at two standard deviations above the mean signal intensity of remote myocardium (2SD), applied to the 6 mm whole-heart MRI dataset. This approach yielded infarct sizes of 47.2 ± 4.0% and 45.5 ± 2.2% of LV volume in WT and L(E) mice, respectively, with no statistically significant difference between genotypes (p = 0.731; **Fig. 6C**). Collectively, these findings demonstrate that the marked attenuation of circulating neutrophilia in L(E) mice was not accompanied by detectable differences in circulating cTnI concentrations or acute myocardial infarct size. These observations suggest that the magnitude of the systemic neutrophilic response following MI does not necessarily predict the extent of acute myocardial injury.

### Longitudinal cardiac MRI reveals comparable ventricular remodelling and systolic dysfunction in WT and L(E) mice following MI

Although acute infarct size is an important determinant of subsequent ventricular remodelling, neutrophils also participate in the inflammatory and reparative processes that influence myocardial healing^5^. We therefore investigated whether the attenuation of post-infarction neutrophilia in L(E) mice influenced the progression of ventricular remodelling following MI. WT and L(E) mice were assessed by longitudinal cardiac MRI at days 1, 14 and 28 following permanent LAD ligation. Late gadolinium-enhanced MRI was used to quantify infarct size, while cine MRI was used to assess LV ejection fraction, end-diastolic volume, end-systolic volume, stroke volume and myocardial mass (**Fig. 7A–E**). Both genotypes exhibited substantial ventricular dysfunction and progressive adverse remodelling over the 28-day observation period.

**Figure 7.**
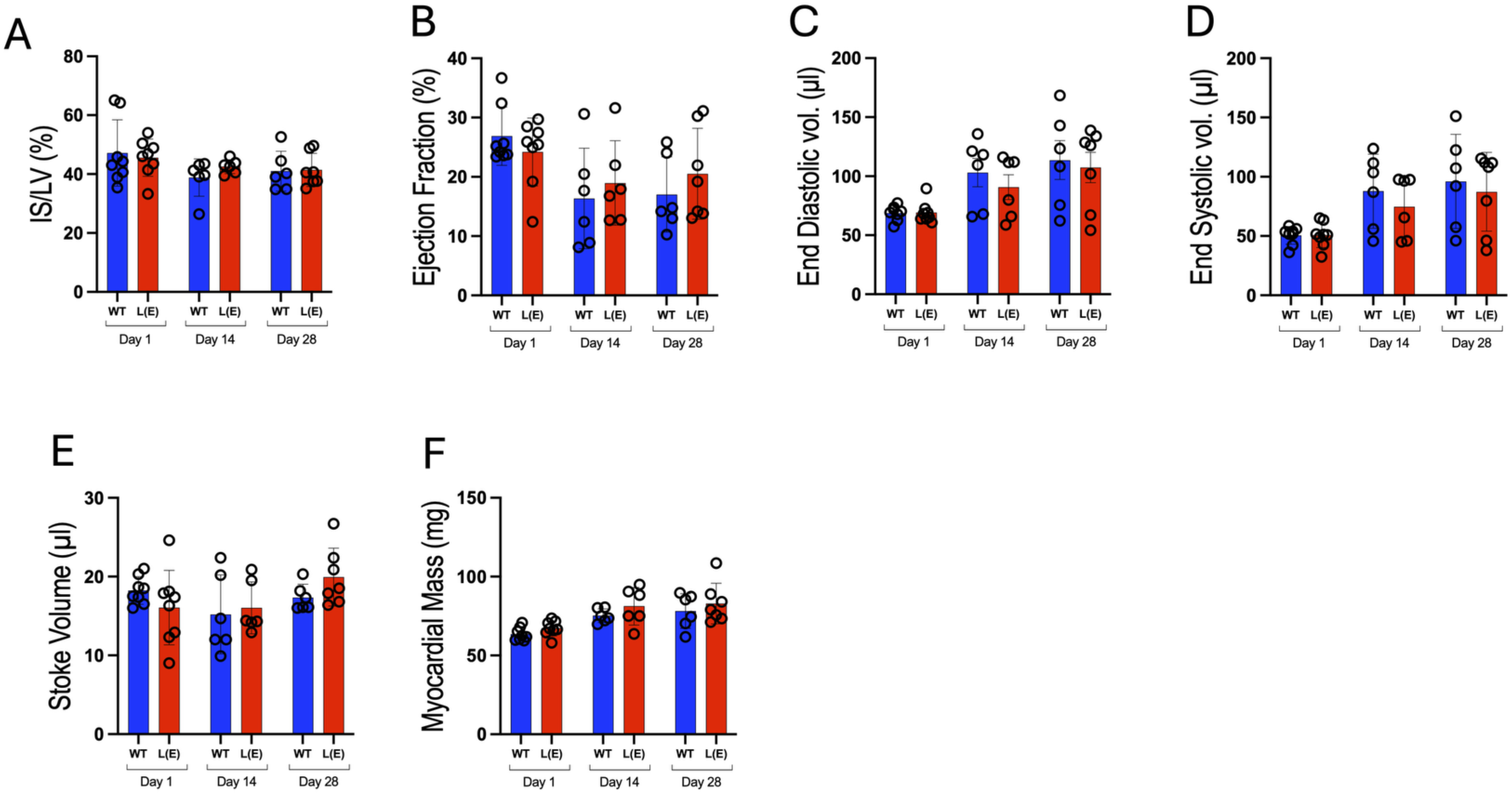
Longitudinal cardiac MRI reveals comparable ventricular remodelling and systolic dysfunction in WT and L(E) mice following myocardial infarction. Wild-type (WT; blue) and L-selectin shedding-resistant (L(E); red) mice underwent permanent ligation of the left anterior descending (LAD) coronary artery and were assessed by cardiac magnetic resonance imaging (MRI) at days 1, 14 and 28 following myocardial infarction (MI). Infarct size was quantified using LGE MRI, applying a semi-automated signal-intensity threshold of two standard deviations above the mean intensity of remote myocardium. Left ventricular (LV) structural and functional parameters were assessed using cine MRI. (**A**) Infarct size (IS/LV %). (**B**) LV ejection fraction (%). (**C**) LV end-diastolic volume (µL). (**D**) LV end-systolic volume (µL). (**E**) LV stroke volume (µL). (**F**) LV myocardial mass (mg). Both genotypes exhibited progressive ventricular dilatation and impaired systolic function following MI, with no statistically significant differences between WT and L(E) mice at any timepoint. Data are presented as mean ± SEM, with each symbol representing an individual mouse (n = 8 per genotype at day 1, n = 6 per genotype at day 14, and n = 6 WT and n = 7 L(E) at day 28). Between-genotype comparisons at each timepoint were performed using unpaired Welch’s t-tests. All comparisons were non-significant (p > 0.05).

At day 14, infarct size was 38.8 ± 2.6% of LV volume in WT mice and 42.6 ± 1.0% in L(E) mice (p = 0.222). By day 28, infarct sizes remained comparable at 41.0 ± 2.8% and 41.4 ± 2.1%, respectively (p = 0.911; **Fig. 7A**). Thus, no statistically significant genotype-associated differences in MRI-derived infarct size were detected throughout the observation period. Next, LV ejection fraction was markedly reduced at day 1 following MI and declined further by day 14 in both genotypes. In WT mice, ejection fraction decreased from 26.9 ± 1.7% at day 1 to 16.4 ± 3.5% at day 14, remaining at 17.0 ± 2.6% at day 28. A similar pattern was observed in L(E) mice, with ejection fractions of 24.2 ± 2.0%, 19.0 ± 2.9% and 20.5 ± 2.9% at days 1, 14 and 28, respectively (**Fig. 7B**). No statistically significant differences were detected between genotypes at any timepoint. Consistent with progressive ventricular dilatation, LV end-diastolic volume increased from 68.8 ± 2.2 µL at day 1 to 113.4 ± 16.4 µL at day 28 in WT mice, compared with an increase from 69.2 ± 3.2 µL to 107.3 ± 12.9 µL in L(E) mice (**Fig. 7C**). Similarly, LV end-systolic volume increased from 50.5 ± 2.6 µL to 96.1 ± 16.2 µL in WT mice and from 50.8 ± 3.7 µL to 87.3 ± 12.6 µL in L(E) mice over the same period (**Fig. 7D**). Despite these changes, neither end-diastolic nor end-systolic volume differed significantly between genotypes at any timepoint. Stroke volume remained broadly comparable between WT and L(E) mice throughout the observation period, with no statistically significant genotype-associated differences at days 1, 14 or 28 (**Fig. 7E**). Likewise, myocardial mass increased over the course of remodelling in both genotypes, without detectable between-genotype differences (**Fig. 7F**).

Together, these findings demonstrate that both WT and L(E) mice develop substantial ventricular dilatation and systolic dysfunction following permanent coronary artery ligation. Despite the pronounced attenuation of circulating neutrophilia in L(E) mice, no detectable differences were observed in the progression of ventricular remodelling, cardiac function or MRI-derived infarct size over 28 days. These observations suggest that attenuation of the systemic neutrophilic response through prevention of L-selectin shedding does not necessarily translate into altered ventricular remodelling following MI.

### L-selectin shedding does not significantly alter myocardial scar formation or structural remodelling at 28 days following MI

To determine whether the altered peripheral neutrophil response observed in L(E) mice was associated with differences in subsequent myocardial repair and remodelling, hearts were examined histologically 28 days following permanent LAD ligation. Picrosirius red staining revealed established collagen-rich infarct scars in both WT and L(E) hearts (**Fig. 8A**).

**Figure 8.**
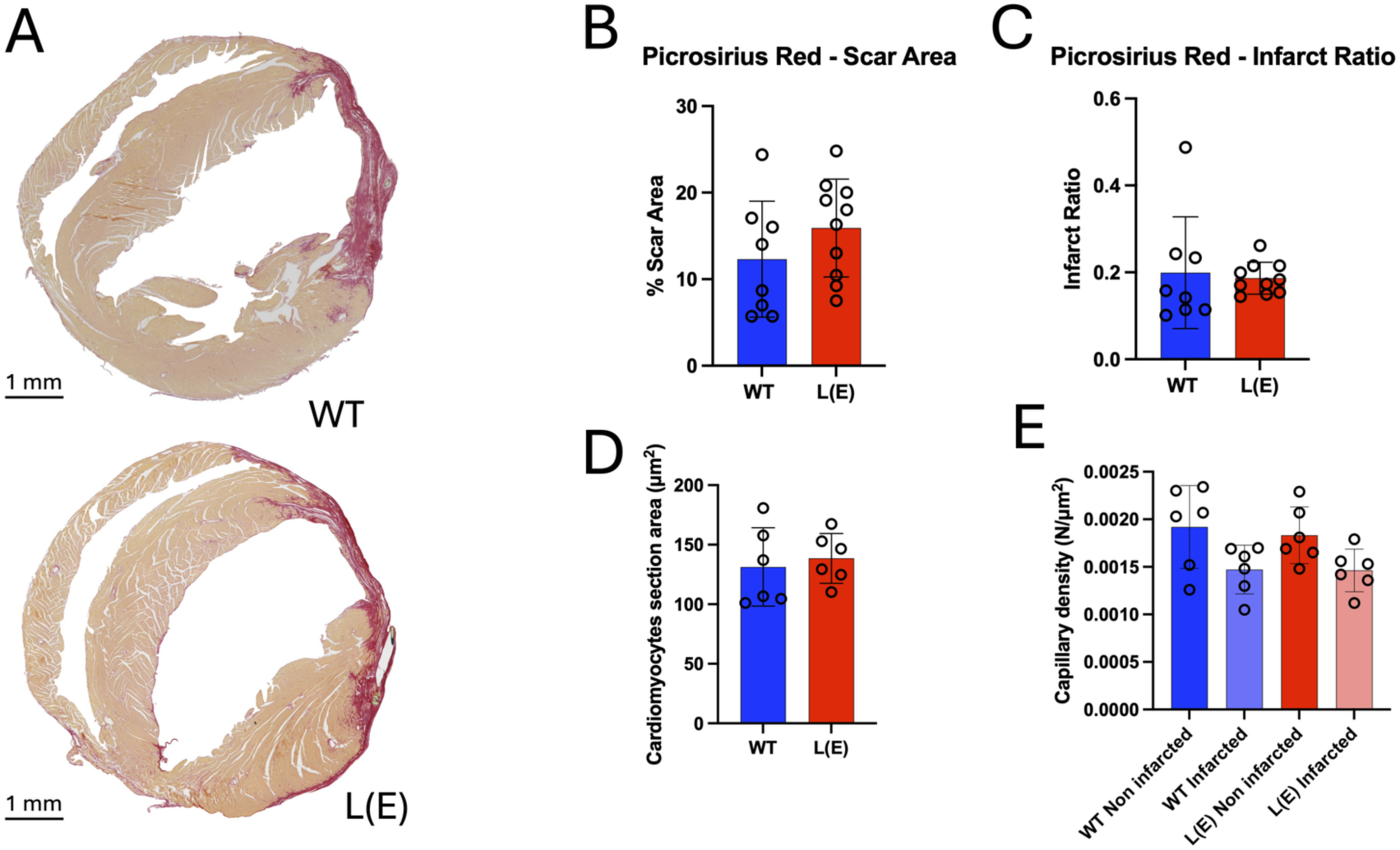
Myocardial scar formation and structural remodelling at 28 days following MI are not significantly altered in L(E) mice. (**A**) Representative short-axis sections of WT and L(E) hearts stained with picrosirius red 28 days following permanent LAD ligation. Collagen-rich scar tissue is stained red, while viable myocardium appears pale orange. Scale bar = 1 mm. (**B**) Myocardial scar area was quantified from picrosirius red-stained sections using colour-threshold-based image analysis in ImageJ/Fiji. Scar area was expressed as a percentage of total myocardial section area. Measurements from four sections per mouse were averaged to obtain a single biological replicate. WT (n = 8) and L(E) (n = 10) mice exhibited comparable scar areas (12.32 ± 6.70% and 15.92 ± 5.65%, respectively; mean ± SD). Between-genotype differences were assessed using a two-tailed unpaired Welch’s t-test (p = 0.245). (**C**) Epicardial infarct ratio, calculated as the infarcted epicardial arc length divided by the total left ventricular epicardial circumference, using an approach adapted from Takagawa et al^30^. WT (n = 8) and L(E) (n = 10) mice exhibited comparable epicardial infarct ratios (19.92 ± 12.85% and 18.66 ± 3.68%, respectively; mean ± SD). Statistical significance was assessed using a two-tailed unpaired Welch’s t-test (p = 0.795). (**D**) Cardiomyocyte cross-sectional area in the peri-infarct border zone, quantified using wheatgerm agglutinin (WGA) staining and semi-automated image segmentation. (**E**) Density of isolectin B4 (IB4)-positive objects in non-infarcted and infarcted myocardial regions, expressed as the number of thresholded IB4-positive objects per µm² of analysed tissue (n = 6 mice per genotype). Paired regional measurements were obtained from each mouse. IB4-positive object density was lower in infarcted than in non-infarcted myocardium in both WT and L(E) mice, with no significant between-genotype differences detected in either region. For panels **B**-**E**, between-genotype comparisons were performed using two-tailed unpaired Welch’s t-tests. Data are presented as mean ± SD, with each biological replicate representing an individual mouse. Blue indicates WT and red indicates L(E) mice. Shaded bars represent the infarcted regions analysed.

Scar formation was quantified using two complementary morphometric approaches. Collagen-positive scar area was expressed as a percentage of total myocardial section area (**Fig. 8B**), while the circumferential extent of infarction was assessed by measuring epicardial infarct length relative to total left ventricular epicardial circumference (**Fig. 8C**), using an approach adapted from Takagawa et al.^30^. Neither measurement revealed a statistically significant difference between WT and L(E) mice.

To investigate additional features of post-infarction remodelling, cardiomyocyte cross-sectional area was assessed in the peri-infarct border zone using wheat germ agglutinin (WGA) staining (**Fig. 8D**). In parallel, isolectin B4 (IB4)-positive object density was quantified in non-infarcted and scarred myocardial regions using semi-automated fluorescence image analysis (**Fig. 8E**). No significant genotype-dependent differences were detected in either endpoint.

Together, these findings indicate that resistance to L-selectin shedding was not associated with detectable changes in myocardial scar extent, border-zone cardiomyocyte cross-sectional area or IB4-positive object density at 28 days following MI.

## Discussion

Here, we show that L-selectin ectodomain shedding contributes to the systemic neutrophilic response following myocardial infarction (MI). Despite normal neutrophil maturation and number under homeostatic conditions, L-selectin shedding-resistant L(E) mice exhibited markedly attenuated circulating neutrophilia following permanent coronary artery ligation. This phenotype was associated with reduced CXCR2 surface expression on mature bone marrow (BM) neutrophils and lower circulating CXCL2 concentrations. Furthermore, recombinant CXCL2 failed to elicit the increase in circulating neutrophil number observed in wild-type (WT) mice, suggesting that reduced chemokine availability alone cannot explain the phenotype. Importantly, attenuated systemic neutrophilia was not accompanied by detectable differences in myocardial neutrophil number, acute myocardial injury or subsequent ventricular remodelling.

The CXCL2/CXCR2 axis is an established regulator of neutrophil mobilisation from the BM, acting in opposition to CXCL12/CXCR4-mediated retention^6–8^. Our findings suggest that L-selectin shedding may influence neutrophil responsiveness to these mobilisation signals. Because neutrophils can themselves produce CXCL2, reduced circulating CXCL2 in L(E) mice could also reflect disruption of an autocrine or paracrine amplification mechanism^9,31^. However, the plasma cytokine-array findings were exploratory and did not remain significant following correction for multiple comparisons. Future studies will be needed to determine the cellular source of circulating CXCL2 and establish the mechanistic relationship between L-selectin shedding and CXCR2 surface expression.

How L-selectin shedding might regulate CXCR2-dependent mobilisation remains an area for further investigation. Retention of L-selectin could alter neutrophil adhesive interactions within the BM microenvironment, while changes in receptor-associated signalling or trafficking might influence CXCR2 surface availability (e.g. influence on receptor recycling or availability of β-arrestin to bind to the tail of CXCR2 for internalisation). We have previously demonstrated that the L-selectin cytoplasmic tail interacts with the μ1A subunit of the AP-1 adaptor complex^32^, and that, in monocytes, ezrin associates with non-cleavable L-selectin during transendothelial migration (TEM)^33^. Whether these interactions contribute to CXCR2 regulation or BM neutrophil mobilisation remains to be established. The timing of neutrophil TEM across inflamed endothelial monolayers is significantly retarded when L-selectin shedding is suppressed^34^. Given the importance of CXCR2 in TEM^31,35^, it is likely that blocking shedding of L-selectin is impacting TEM through similar disrupted CXCR2 signalling/sensing mechanisms observed in this study.

Our findings also complement those of Anzai et al.^36^, who demonstrated that cardiac fibroblast-derived GM-CSF promotes inflammatory myeloid cell responses following MI. Although circulating GM-CSF and G-CSF were not detectably altered in L(E) mice (**Fig. 4A**), reduced CXCR2 expression on mature BM neutrophils suggests that L-selectin shedding may influence chemokine responsiveness independently of changes in circulating growth-factor concentrations. Nevertheless, interactions between these pathways cannot be excluded.

Perhaps the most interesting observation is that the pronounced attenuation of circulating neutrophilia in L(E) mice was not accompanied by a detectable reduction in the relative number of neutrophils within the infarcted myocardium. Recent evidence suggests that vascular marginated neutrophil populations contribute substantially to early (6h) myocardial recruitment following MI^1^. Our findings are compatible with the possibility that myocardial recruitment can be sustained without a proportionate increase in newly mobilised BM neutrophils.

The relationship between neutrophil mobilisation and cardiac outcome is similarly complex. Previous studies have demonstrated that neutrophils contribute not only to inflammatory injury but also to subsequent myocardial repair. Neutrophil depletion can impair reparative macrophage responses and ventricular recovery^5^, whereas disruption of GM-CSF-dependent inflammatory cell mobilisation has been associated with improved post-infarction outcomes^36^.

Furthermore, circadian variation in neutrophil recruitment to the infarcted myocardium has been shown to influence infarct severity and subsequent cardiac repair^37^. In contrast, the attenuated circulating neutrophilic response in L(E) mice was not associated with detectable differences in acute injury, longitudinal ventricular function or chronic scar formation. These findings suggest that the magnitude of systemic neutrophilia alone may not determine cardiac outcome following permanent coronary occlusion.

In conclusion, our findings identify L-selectin shedding as a regulator of the systemic neutrophilic response following MI, potentially through mechanisms involving CXCL2/CXCR2 signalling. The dissociation between attenuated circulating neutrophilia and the absence of detectable changes in myocardial injury or remodelling highlights the importance of distinguishing systemic neutrophil mobilisation from local inflammatory responses within the infarcted heart.

## Materials and methods

### Animals and genotyping

#### Ethical statement

Experiments used mice carrying a knock-in allele encoding a cleavage-resistant form of L-selectin, in which the membrane-proximal ADAM17 cleavage site is replaced by the equivalent sequence of E-selectin^15^, hereafter L(E), together with their wild-type (WT) littermates. The line was a gift from Professor Douglas Steeber (University of Wisconsin-Milwaukee) and was maintained on a C57BL/6 background. Mice were group-housed on a 12 h light/dark cycle with food and water available at all times. All mice used in this report were aged 8-13 weeks (MI surgery, systemic CXCL2 challenge, peritonitis model, and ex vivo stimulations). All procedures were approved in advance by the King’s College London Ethical Review Process Committee and performed under UK Home Office Project Licences 70/8198, 70/8889, 70/7959 and PA93DA655. Work complied with the Animals (Scientific Procedures) Act 1986, European Directive 2010/63/EU and NC3Rs guidance, and is reported in accordance with the ARRIVE guidelines.

#### Genotyping

Genotypes were assigned by PCR followed by agarose gel electrophoresis. Ear-clip biopsies were digested by boiling in 300 µL of 50 mM NaOH for 30 min, after which 50 µL of 1 M Trizma base, pH 8.0, was added to neutralise the lysate. Two microlitres of this crude extract served as template. Reactions were assembled in a final volume of 25 µL using GoTaq Green Master Mix (Promega) with each of four primers at 250 nM (sequences in Table 1). Hot-start cycling on a Veriti thermal cycler (Thermo Fisher Scientific, UK) comprised an initial denaturation of 7 min at 94 °C; 35 cycles of 94 °C for 45 s, 56 °C for 45 s and 72 °C for 45 s; and a terminal extension of 3 min at 72 °C, after which reactions were held at 4 °C. Products were resolved on a 2.5% (w/v) agarose gel cast in Tris-Acetate-EDTA (TAE) buffer containing 5000-fold dilution (v/v) Nancy-520 (Sigma-Aldrich, UK), run at 150 V for 35 min, and visualised under ultraviolet illumination on a ChemiDoc imaging system (Bio-Rad Laboratories). The wild-type allele yields a 349 bp product and the L(E) allele a 202 bp product. See Table 1 for PCR primer designs.

**Table 1.**
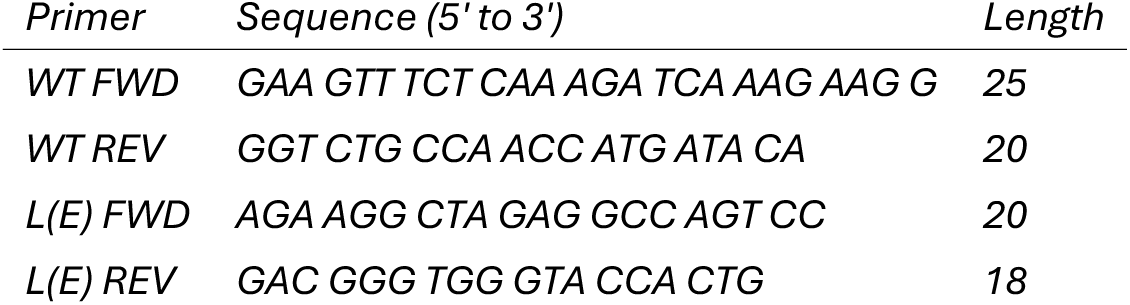
Genotyping primers. All primers from Integrated DNA Technologies, US.

#### Sex as a biological variable

Neutrophilia and myocardial neutrophil recruitment occur after infarction in mice of both sexes, although their magnitude is greater in males^38–40^. In this study, infarction studies were performed exclusively in female mice. This was a deliberate choice made on welfare grounds: male mice subjected to permanent coronary ligation show a higher incidence of left ventricular free wall rupture in the days following infarction^38^, and restricting the surgical cohort to females was adopted to reduce unplanned mortality and the number of animals required, in keeping with the 3Rs. Neutrophilia is a conserved feature of the acute response to infarction in mice of both sexes, and neutrophil accumulation within the infarct peaks at 24 h in males and females alike^40^. The systemic CXCL2 challenge, which involves no surgery and carries no comparable mortality risk, was performed in male mice, and provides the only in vivo assessment of neutrophil mobilisation in male animals. Because the infarction and chemokine challenge experiments differ in both stimulus and sex, they do not constitute a formal test of sex as a biological variable, and no statistical comparison between sexes was possible.

#### Myocardial infarction surgery

Infarction was produced by permanent occlusion of the left anterior descending (LAD) coronary artery. Female mice aged 8-13 weeks received ketamine (75 mg/kg; Vetalar, Bioniche Animal Health) and medetomidine hydrochloride (1 mg/kg; Domitor, Orion Pharmaceuticals) by intraperitoneal injection, together with buprenorphine (0.15 mg/kg, intramuscular; Vetergesic, Alstoe Animal Health, UK) and flunixin meglumine (2.5 mg/kg, subcutaneous; Finadyne, Schering-Plough) for perioperative analgesia. The chest was shaved, and animals were intubated and ventilated at 150 breaths/min with a tidal volume of 150 µL (MiniVent 845, Hugo Sachs/Harvard Apparatus, UK).

The heart was exposed through a lateral thoracotomy at the fourth intercostal space. The LAD was ligated permanently 1-2 mm distal to the left atrium with an 8/0 polypropylene suture (Ethicon, Johnson & Johnson Medical, UK). Occlusion was considered successful when blanching extended from the suture towards the apex. The chest was closed, anaesthesia reversed with atipamezole (5 mg/kg, intraperitoneal; Antisedan, Pfizer), and animals recovered in a heated chamber. Sham-operated mice underwent an identical procedure in which the suture was passed beneath the LAD but not tied. All surgery was performed by experienced microsurgeons blinded to genotype.

#### Heart harvest and preparation for immunohistochemistry

At 24 h after surgery mice were sacrificed by a Schedule 1 method. The diaphragm and sternum were opened to expose the heart, which was excised through the aorta and venae cavae; thymic tissue was carefully removed to avoid contaminating subsequent leukocyte preparations. Hearts were transferred while still beating into ice-cold PBS containing 5% KCL (v/v) to arrest in diastole -prior to fixation in 4% PFA (overnight at 4°C) and subsequent paraffin embedding. Each heart was then divided in the transverse plane at the level of the ligature, separating the remote and non-infarcted base from the infarcted apex.

#### Heart collection for cardiac leukocyte isolation

At 24 h following myocardial infarction or sham surgery, mice were euthanised using an approved Schedule 1 method. Hearts were excised following opening of the thoracic cavity, and surrounding thymic tissue was carefully removed. Excised hearts were transferred immediately into ice-cold PBS to facilitate clearance of residual intravascular blood. The site of coronary artery ligation, or the corresponding anatomical position in sham-operated mice, was identified, and hearts were divided transversely into basal and apical regions. Tissue from each region was subsequently processed separately for enzymatic digestion and flow-cytometric analysis.

#### Collagenase digestion and retrieval of cardiac-resident leukocytes

Base and apex fragments were minced and digested in 300 µL of PBS containing DNase I (400 pg/mL; bovine pancreas, type IV, Sigma-Aldrich, UK), hyaluronidase type I-S (500 U/mL) (bovine testes, Sigma-Aldrich, UK) and collagenase IV (1 mg/mL; Clostridium histolyticum, Sigma-Aldrich, UK) for 30 min at 37 °C in 5% CO2. Digestion was arrested by adding EDTA and EGTA in PBS to 5 mM each, chelating the divalent cations required for enzyme activity. The suspension was filtered through a 40 µm strainer into 5 mL of ice-cold calcium- and magnesium-free HBSS, centrifuged at 450 g for 5 min and resuspended in complete medium.

#### Antigen retrieval

Hearts were fixed overnight at 4 °C in 4% (w/v) paraformaldehyde before paraffin embedding and sectioning. Sections were dewaxed in two 5 min changes of 100% xylene followed by 5 min in a 1:1 mixture of xylene and absolute ethanol, and rehydrated through two 5 min changes each of 100% and 90% ethanol, two 3 min changes of 70% ethanol and two 5 min changes of distilled water.

Heat-induced epitope retrieval was performed in 10 mM Tris base, 1 mM EDTA, pH 9.0. The buffer was pre-heated in a staining jar to 95–100 °C in a steamer, slides were immersed, the lid placed loosely on the jar, and sections incubated for 30 min. Slides were then allowed to cool for at least 30 min at room temperature, washed twice for 3 min in PBS, and a hydrophobic barrier drawn around each section (DAKO pen S2002, Agilent Technologies).

Sections were permeabilised in 0.2% (v/v) Triton X-100 for 5 min at 4 °C and washed three times for 3 min in PBS. Blocking was performed for 60 min at room temperature in a humidified chamber in PBS containing 20% (v/v) donkey serum, 0.05% (v/v) Tween-20 and 0.6 M glycine. Blocking solution was removed by flicking without a subsequent wash, and sections were incubated overnight at 4 °C in a humidified chamber with isolectin B4-Alexa Fluor 488 (1:200, 1 mg/mL; ThermoFisher), WGA-TRITC (1:200, 1 mg/mL; ThermoFisher), together with DAPI at 1 ng/mL (ThermoFisher), all diluted in PBS containing 0.05% (v/v) Tween-20 and 0.6 M glycine. Sections were washed three times for 3 min in PBS/Tween-20 and once for 3 min in distilled water. Lipofuscin autofluorescence was quenched by incubation in 0.1% (w/v) Sudan Black B in 70% ethanol for 2-3 min at room temperature, followed by washing in running tap water until the sections cleared and a final 3 min wash in distilled water. Sections were mounted in Fluoromount-G (SouthernBiotech) and stored at 4 °C.

#### CXCL2-induced peritoneal neutrophil recruitment

Mice received an intraperitoneal injection of recombinant murine CXCL2 (R&D Systems; 0.5 µg in 100 µL PBS). Four hours after administration, animals were euthanised by cervical dislocation, and peritoneal leukocytes were recovered by lavage. Following exposure of the peritoneal cavity, 5 mL of ice-cold PBS was introduced using a 10 mL syringe fitted with a 19-gauge needle. The abdomen was gently massaged to facilitate cell recovery before the lavage fluid was aspirated. This procedure was repeated once, yielding approximately 10 mL of pooled lavage fluid per animal. Recovered cells were centrifuged at 450 × g for 5 min, subjected to erythrocyte lysis and washed before resuspension in 2% (v/v) BSA in PBS. Cells were subsequently stained with the appropriate fluorochrome-conjugated antibodies containing FcR Blocking Reagent (Miltenyi Biotec) and analysed by flow cytometry, as described in the relevant Methods subsection.

#### Cardiac magnetic resonance imaging

Female mice underwent cardiac MRI at baseline and/or at 1, 14 and 28 days following permanent LAD ligation, according to the experimental protocol. Gadobutrol (0.65 mmol/kg per mouse; Gadovist, Bayer Pharmaceuticals) was administered intraperitoneally as a contrast agent for late gadolinium enhancement, and imaging began approximately 45 minutes after injection. Anaesthesia was induced with 5% and maintained with 2% (v/v) isoflurane (Isoflo, Abbott Laboratories). Core temperature was maintained using a water-filled heating blanket, and temperature and respiratory rate were monitored throughout.

Imaging was performed on a 9.4 T small-animal MRI system (Biospec 94/20, Bruker Biospin, DE) with a 39-mm quadrature volume coil (RAPID Biomedical, DE). Cardiac cine MRI was performed using a retrospectively gated gradient-echo sequence (IgFLASH; Bruker) with six consecutive short-axis slices covering approximately 6 mm of the left ventricle. Each slice was 1 mm thick, with a 30 × 30 mm field of view and a 192 × 192 acquisition matrix, providing an in-plane resolution of approximately 156 µm. The repetition time and echo time were 8.262 and 1.8775 ms, respectively, and ten cardiac phases were reconstructed per slice.

Images were analysed in cvi42 (version 5.13; Circle Cardiovascular Imaging Inc.) by a single operator blinded to genotype. For functional analysis, endocardial and epicardial contours were generated using the automated detection tool and inspected on every short-axis cine slice, with manual correction where the automated contour did not follow the myocardial border. End-diastolic and end-systolic phases were taken as the frames of largest and smallest left ventricular volume.

Myocardial infarct size was quantified from LGE short-axis images using a semi-automated signal-intensity threshold method applied to the six-slice short-axis LV analysis dataset. Endocardial and epicardial contours were delineated to define the myocardial boundaries, and a region of interest was placed within remote, non-enhancing myocardium to establish the reference signal intensity. Infarcted myocardium was identified as tissue exhibiting signal intensity greater than two standard deviations (2SD) above the mean signal intensity of the remote reference region. The resulting segmentation was reviewed for inclusion of non-myocardial signal or imaging artefacts. The same thresholding approach was applied to WT and L(E) datasets. Infarct size was expressed as the percentage of total LV volume occupied by LGE-positive myocardium (IS/LV%).

### Cell and tissue isolation

#### Blood and plasma

Mice were terminally anaesthetised with 3% (v/v) isoflurane and exsanguinated by cardiac puncture. With the animal supine, a 23G needle was advanced through the diaphragm at the midline into the apex of the left ventricle and up to 1 mL of blood withdrawn directly into a tube containing 50 U heparin (B. Braun Medical, UK). Samples were held on ice. Death was confirmed by cervical dislocation. For plasma, heparinised blood was centrifuged at 600 g for 15 min at 4 °C and the supernatant collected and stored at −80 °C.

Leukocytes were recovered from 200 µL of whole blood by lysing erythrocytes in 10 mL of ammonium chloride haemolysis buffer (155 mM NH_4_Cl, 10 mM KHCO_3_, 1 mM EDTA). Cells were pelleted at 450 *g*, washed once in PBS, pelleted again and resuspended in complete medium (RPMI 1640 with 5 mM L-glutamine (Gibco, Thermo Fisher Scientific, UK), 10% (v/v) heat-inactivated foetal bovine serum (Sigma-Aldrich, UK) and 1% (v/v) penicillin-streptomycin (Thermo Fisher Scientific, UK)).

#### Bone marrow isolation

After Schedule 1 killing, hind limbs were removed by blunt dissection and the femur and tibia isolated. Marrow was flushed from each bone with calcium- and magnesium-free HBSS (Sigma-Aldrich, UK) through a 27G needle, passed through a 40 µm strainer and resuspended thoroughly to give a single-cell suspension. Cells were pelleted at 450 *g*, subjected to haemolysis in 10 mL of ammonium chloride buffer, pelleted and washed once in PBS. For direct analysis by flow cytometry, cells were resuspended in complete medium and kept on ice until use. Where purified neutrophils were required, the washed pellet was instead resuspended in MACS buffer and taken directly into magnetic separation as described below.

#### Isolation of neutrophils from bone marrow

Neutrophils were isolated by negative magnetic selection using the Neutrophil Isolation Kit, mouse (Miltenyi Biotec; 130-097-658) according to the manufacturer’s instructions. Non-target cells were labelled with a cocktail of biotin-conjugated antibodies directed against antigens absent from neutrophil granulocytes, followed by anti-biotin MicroBeads, and retained on an LS column (Miltenyi Biotec) in the field of a MidiMACS separator; unlabelled neutrophils were collected in the flow-through, so that the isolated cells were untouched by antibody. All steps were performed at 2-8 °C in degassed MACS buffer (PBS, pH 7.2, containing 0.5% (w/v) BSA and 2 mM EDTA), and cells were passed through a 30 µm pre-separation filter before loading. Purity was assessed by flow cytometry as the proportion of Ly6G⁺ events and was routinely ≥95%. Isolated neutrophils were washed to remove EDTA-containing buffer and resuspended in the appropriate medium for subsequent ex vivo stimulation assays.

### Flow cytometry

#### Staining and acquisition

Antibody cocktails were made up in PBS containing 5% (v/v) rat serum (Thermo Fisher Scientific, UK), each antibody at a final concentration of 2 µg/mL. Cells were resuspended in 100 µL of staining solution and held on ice in the dark for at least 90 min. Unbound antibody was diluted out with ice-cold calcium- and magnesium-free HBSS, cells were centrifuged at 450 g and fixed in 1% (v/v) paraformaldehyde in PBS. Fixed samples were stored at 4 °C in the dark until acquisition.

Data were acquired on LSRFortessa or FACSSymphony cytometers (BD Biosciences) at the NIHR Biomedical Research Centre, King’s College London and Guy’s and St Thomas’ NHS Foundation Trust, or at the MRC Centre for Transplantation, King’s College London. Compensation was set before each run using single-stained UltraComp eBeads (Invitrogen, Thermo Fisher Scientific, US). Acquisition used FACSDiva v8 (BD Biosciences) and analysis FlowJo v10 (FlowJo LLC). Unstained and fluorescence-minus-one controls defined background autofluorescence.

#### Neutrophil identification in blood and bone marrow

Single cells were gated on forward and side scatter, doublets excluded on SSC-W against SSC-A, and CD45+ leukocytes selected. Neutrophils were identified within this gate as SSC-int Ly6G+ events. A stopping gate of 30,000 CD45+ leukocytes was collected per sample and neutrophil number expressed per 30,000 CD45+ cells. Surface expression of L-selectin, Mac-1 (CD11b), CXCR2 and CXCR4 was recorded as median fluorescence intensity. The panel comprised CD45-APC, Ly6G-FITC, CD11b-BV510 and L-selectin-PE (all from BD Biosciences).

#### Neutrophil identification in digested heart (supplemental Figure 1)

Cardiac leukocytes were stained with CD45-APC-Cy7, Ly6C-BV605, Ly6G-BV785, L-selectin-PE and CD11b-BV510. Neutrophils were defined as CD45+Ly6C-int Ly6G+ events within a stopping gate of 30,000 CD45+ leukocytes, and L-selectin and Mac-1 expression recorded as median fluorescence intensity.

#### Bone marrow neutrophil maturation

Maturation stage was assigned using the published gating scheme of Evrard et al. (2018)^24^, with antibodies against CD170-PE, CD115-PE-Cy7, Gr1-FITC, CD11b-BV510, CD117-BV711, CXCR4-APC, CXCR2-BV421 and Ly6G-BV785. After exclusion of CD115+ monocytes and CD170+ eosinophils, Gr1+CD11b+ neutrophils were gated and three subsets resolved within it: CD117-hi CXCR4-hi pre-neutrophils, Ly6G-lo CXCR2-lo immature neutrophils, and Ly6G+CXCR2+ mature neutrophils. Each subset is expressed as a percentage of total neutrophils, and CD11b, CXCR2 and CXCR4 expression was measured as median fluorescence intensity within each. A stopping gate of 10,000 CD170-lo CD115-lo Gr1+CD11b+ neutrophils was collected per sample.

#### Chemoattractant stimulation ex vivo

Leukocytes from blood or bone marrow were brought to 10 × 10⁶ cells/mL in complete medium and 50 µL aliquots (5 × 10⁵ cells) dispensed into U-bottom 96-well plates (Nunclon Delta Surface, Thermo Fisher Scientific, UK). Where metalloproteinase inhibition was required, suspensions received TAPI-0 (Santa Cruz Biotechnology; sc-203410) at 10 µM and were incubated for 30 min at 37 °C in 5% CO2 before stimulation; vehicle controls received an equal volume of DMSO and the same pre-incubation period.

Chemoattractants were prepared at twice the working concentration (1 µM) in complete medium and 50 µL added to each well, halving the concentration to 500 nM final and the cell density to 5 × 10⁶/mL. Where cells had been pre-treated, TAPI-0 or DMSO was carried into the chemoattractant preparation at the same concentration. The agonists used were recombinant murine CXCL2/MIP-2 (R&D Systems, Bio-Techne; 452-M2/CF), leukotriene B4 (Sigma-Aldrich, UK; L0517) and the formyl peptide WKYMVm (Tocris, Bio-Techne; 1800). Plates were incubated for 30 min at 37 °C in 5% CO_2_, after which stimulation was stopped by adding 200 µL of ice-cold calcium- and magnesium-free HBSS per well. Cells were then stained for flow cytometry with CD45-APC, Ly6G-FITC, CD11b-BV510 and L-selectin-PE as above.

Surface L-selectin and Mac-1 are reported as median fluorescence intensity, normalised where indicated to unstimulated cells of the same genotype.

#### Protein microarray and statistical analysis

Plasma chemokine and cytokine profiling was performed using a RayBio G-Series mouse cytokine antibody array (RayBiotech, Peachtree Corners, GA, USA; G-series AAM-CYT-G3/G4/G5 for arrays 3, 4 and 5), which together detect 144 targets comprising cytokines, chemokines, growth factors, soluble receptors, adhesion molecules and proteases. Each target is printed in duplicate within each sub-array, and each glass slide carries multiple identical sub-arrays allowing several samples to be processed in parallel. Plasma was collected from WT and L(E) mice at days 1 and 5 after permanent LAD ligation (n = 4 mice per genotype per timepoint; the day 1 and day 5 samples were obtained from independent cohorts of animals). Slides were equilibrated to room temperature, air dried, and blocked with sample diluent for 30 min. Samples were applied at 100 µl per sub-array following 5-fold dilution in sample diluent and incubated overnight at 4°C. Sub-arrays were washed five times in wash buffer I and twice in wash buffer II, incubated with the biotinylated detection antibody cocktail for 2h, washed as before, and incubated with Cy3-equivalent dye-conjugated streptavidin for 1 h in the dark. Slides were washed, dried, and scanned on an InnoScan 710 IVD laser scanner (Innopsys, Carbonne, France) at a Cy3 excitation wavelength, with spot intensities extracted using Mapix software. Day 1 and day 5 sample sets were processed and scanned as two separate batches.

For each spot, the median foreground fluorescence intensity was corrected by subtraction of the local background, and replicate spots were averaged. Signals were then background-corrected using the negative control spots and normalised to the positive control spots of each sub-array, scaling each sub-array to the mean positive control signal across the slide as recommended by the manufacturer. Normalised intensities were log2-transformed for all analyses. Array signal is multiplicative and approximately log-normally distributed, with a near-constant coefficient of variation across groups; transformation stabilised the variance and permitted the use of parametric tests (Levene’s test on log2 values, p = 0.66 and p = 0.60 at days 1 and 5 respectively, against a 19.7-fold between-genotype variance ratio on the raw scale at day 1). For the array-wide survey, the effect of genotype was estimated by two-way ANOVA with genotype and day as fixed factors; because the day 1 and day 5 samples derive from different animals, no repeated-measures matching was applied across timepoints. P values were corrected for testing of 144 analytes by the Benjamini–Hochberg procedure. No analyte reached a false discovery rate below 0.05, and the array was therefore used to identify candidates for direct testing rather than as a source of statistical inference. Plasma CXCL2 was subsequently compared between genotypes at each timepoint by unpaired Student’s t-test on log2-transformed intensities, with the day 1 and day 5 cohorts analysed independently and no correction applied across the two timepoints. Rank-based tests were not used because, with n = 4 mice per genotype per day, the smallest attainable two-sided p value is 0.029 irrespective of effect size.

#### Systemic CXCL2 challenge

To test whether exogenous chemokine could drive neutrophil release from marrow, male mice aged 8-12 weeks received 1 µg of recombinant murine CXCL2 (R&D Systems, Bio-Techne; 452-M2/CF) in 100 µL of sterile PBS by tail vein injection; controls received 100 µL of PBS alone. Blood was collected 45 min after injection by cardiac puncture and leukocytes isolated as described above. Cells were stained with CD45-APC and Ly6G-BV785 and circulating neutrophils enumerated by flow cytometry within a stopping gate of 30,000 CD45+ leukocytes.

#### Histological assessment and quantification of myocardial scar formation

Hearts were collected 28 days following permanent ligation of the left anterior descending (LAD) coronary artery. Short-axis myocardial sections were stained with picrosirius red to visualise collagen deposition and imaged using wide-field microscopy. Digital images were analysed using ImageJ/Fiji with a custom macro (PicrosiriusRedMacro_2024.ijm) employing hue, saturation and brightness (HSB)-based colour thresholding.

For each section, total tissue area was identified using HSB thresholds of 0–255 for hue, 0–255 for saturation and 1–254 for brightness. Collagen-positive scar tissue was identified separately using thresholds of 40–255 for hue, 60–255 for saturation and 0–254 for brightness. Binary masks were generated from the thresholded images, and the corresponding areas were quantified. The resulting segmentations were visually inspected.

Scar area was expressed as a percentage of total tissue-section area, calculated by dividing collagen-positive scar area by total section area and multiplying by 100. In addition, the circumferential extent of infarction was assessed using a length-based approach adapted from Takagawa et al.^30^. Epicardial infarct length was measured along the scarred region of the left ventricular epicardial boundary and normalised to the total epicardial circumference, yielding a dimensionless epicardial infarct ratio.

Four short-axis sections were analysed per animal, and section-level measurements were averaged to generate a single biological replicate per mouse. Histological measurements were compared between wild-type (WT) and L-selectin shedding-resistant (L(E)) mice.

#### Cardiomyocyte cross-sectional area

Fluorescence-stained myocardial sections were imaged using a Hamamatsu NanoZoomer whole-slide scanner at 40× magnification. Images were processed using custom ImageJ/Fiji macros incorporating contrast-limited adaptive histogram equalisation (CLAHE) and background subtraction. Wheat germ agglutinin (WGA) staining was used to delineate cardiomyocyte boundaries within manually selected regions of the peri-infarct border zone. The red fluorescence channel was isolated, thresholded between 0 and 60 and converted to a binary mask. Segmented cardiomyocyte profiles were quantified using the ImageJ/Fiji Analyze Particles function, with a minimum particle area of 30 calibrated area units, circularity of 0.30–1.00 and exclusion of particles touching image boundaries. Mean profile area was expressed in µm². Measurements from four myocardial sections were aggregated to generate one biological replicate per mouse (n = 6 per genotype).

#### Quantification of IB4-positive objects

Isolectin B4 (IB4)-stained myocardial sections were imaged using a Hamamatsu NanoZoomer whole-slide scanner at 40× magnification and analysed using custom ImageJ/Fiji macros. Regions corresponding to non-infarcted myocardium and infarct scar were selected separately. For each region, tissue area was determined from the blue fluorescence channel following thresholding and binary-mask processing. IB4-associated fluorescence was quantified from the green channel using a threshold of 22–255, followed by binary conversion and particle analysis with a minimum particle area of 10 calibrated area units and circularity of 0.00–1.00. IB4-positive object density was calculated as the number of detected fluorescent particles divided by the corresponding analysed tissue area (µm²). This measurement represents the density of thresholded IB4-positive objects rather than a manually validated count of individual capillaries. Measurements from four myocardial sections were aggregated to generate one biological replicate per mouse (n = 6 per genotype).

#### Statistical analysis

Statistical analyses and graphical representations were performed using GraphPad Prism version 10 (GraphPad Software), with descriptive statistics additionally calculated using Microsoft Excel version 16 (Microsoft). Data are presented as mean ± standard deviation (SD) or mean ± standard error of the mean (SEM), as specified in the corresponding figure legends. The biological replicate was the individual mouse for in vivo experiments and an independent experimental preparation for in vitro assays. For histological analyses, measurements from four myocardial sections were aggregated to generate a single value per mouse, and statistical comparisons were performed using these mouse-level values.

Comparisons between two independent groups were performed using two-tailed unpaired Student’s t-tests or Welch’s t-tests, as appropriate. For experiments involving more than two groups, ordinary one-way analysis of variance (ANOVA) or Welch’s ANOVA was performed, followed by Dunnett’s or Dunnett’s T3 multiple comparisons test, respectively. Experiments involving two independent variables were analysed using two-way ANOVA followed by Šidák’s multiple comparisons test, as indicated in the figure legends.

For the plasma cytokine antibody array, signal intensities were log₂-transformed before analysis by two-way ANOVA, with genotype and post-infarction timepoint included as factors.

To account for multiple testing across 144 analytes, p values were adjusted using the Benjamini–Hochberg false discovery rate procedure. Individual CXCL2 comparisons between genotypes at each timepoint were performed using unpaired Student’s t-tests on log₂-transformed intensities and were considered exploratory.

For cardiac MRI, comparisons between WT and L(E) mice at individual timepoints were performed using unpaired Welch’s t-tests. The same animals contributed to the day-1 MRI analyses presented in Figs 6 and 7; these measurements were not treated as independent biological cohorts.

All statistical tests were two-sided, with statistical significance defined as p < 0.05, except where multiplicity-adjusted significance thresholds were applied. Sample sizes and the statistical tests used for individual datasets are specified in the corresponding figure legends.

## Acknowledgements

This work was generously supported by the British Heart Foundation Project Grant: PG/19/42/34429 to Davide Maselli. Samuel Hart was generously supported by the BHF Centre of Research Excellence studentship at King’s College London (RE/13/2/30182). Yue Yang was generously supported by the Chinese Scholarship Council, in collaboration with King’s College London.

---

**Supplementary Figure 1.**
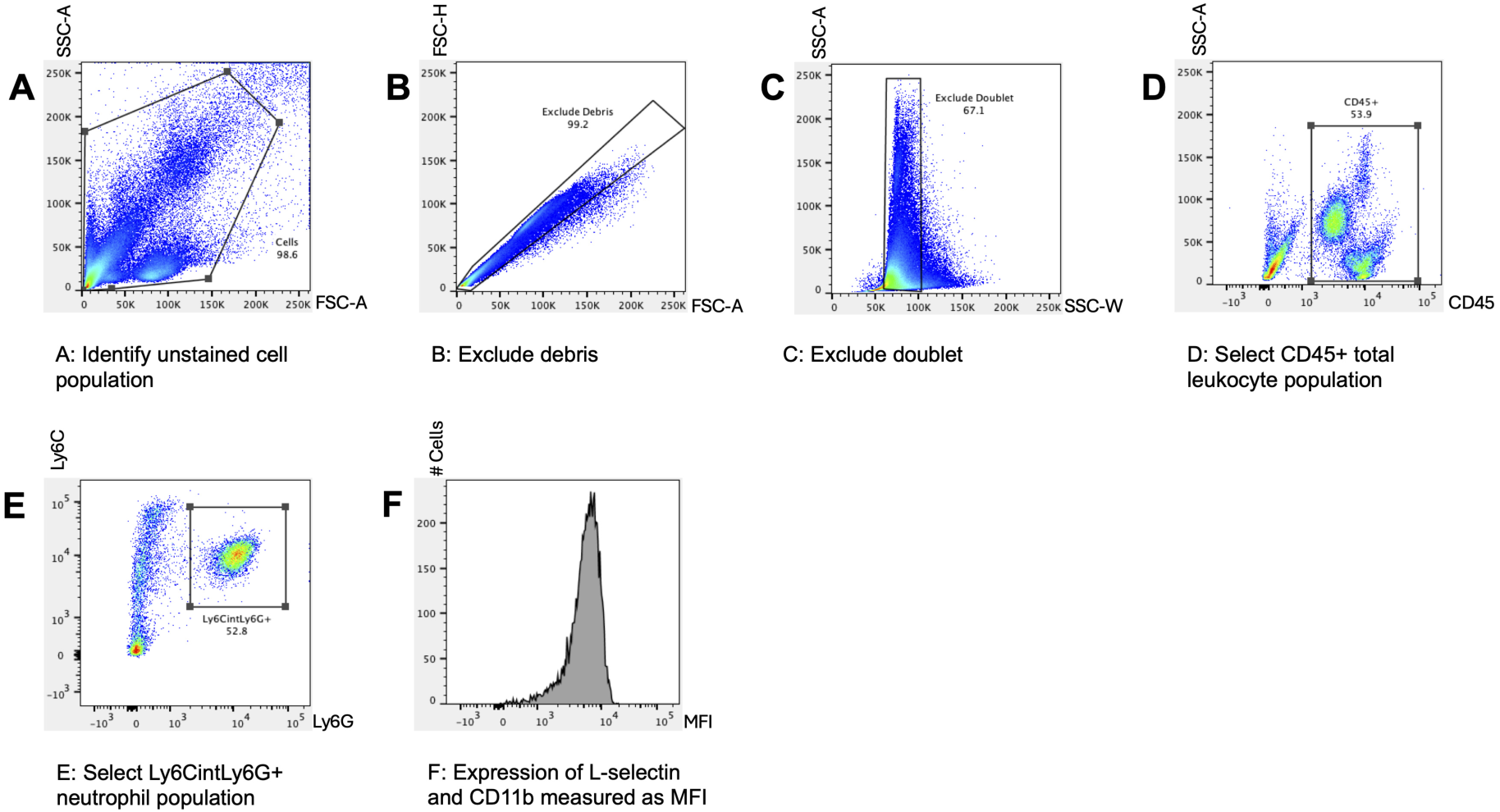
Flow-cytometric identification of neutrophils isolated from cardiac tissue. Representative gating strategy for the identification of CD45⁺ Ly6C^int^ Ly6G⁺ neutrophils in enzymatically digested mouse myocardium. (A) The cellular population was identified using forward-and side-scatter characteristics. (B) Debris was excluded using forward-scatter parameters. (C) Doublets were excluded using side-scatter area and width. (D) CD45⁺ leukocytes were selected. (E) Neutrophils were identified as Ly6C^int^ Ly6G⁺ events within the CD45⁺ population. (F) Surface expression of L-selectin and CD11b was quantified as median fluorescence intensity (MFI) within the gated neutrophil population. A fixed acquisition population of 30,000 CD45⁺ leukocytes was used for the cardiac analyses.

